# A blood-based prognostic biomarker in inflammatory bowel disease

**DOI:** 10.1101/535153

**Authors:** D. Biasci, J.C. Lee, N.M. Noor, D.R. Pombal, N Lewis, T Ahmad, A Hart, M Parkes, E.F. McKinney, P.A. Lyons, K.G.C. Smith

## Abstract

**Objective:** We have previously described a prognostic transcriptional signature in CD8 T cells that separates inflammatory bowel disease (IBD) patients into 2 phenotypically-distinct subgroups, which we termed IBD1 and IBD2. Here we sought to develop a blood-based prognostic test that could identify these patient subgroups without the need for cell separation, and thus be suitable for routine clinical use in Crohn’s disease (CD) and ulcerative colitis (UC).

**Design:** Patients with active IBD were recruited before treatment. Transcriptomic analyses were performed on purified CD8 T cells and/or whole blood. Detailed phenotype information was collected prospectively. IBD1/IBD2 patient subgroups were identified by consensus clustering of CD8 T cell transcriptomes. In a training cohort, statistical (machine) learning was used to identify groups of genes (“classifiers”) whose differential expression in whole blood re-created the IBD1/IBD2 patient subgroups. Genes from the best classifiers were qPCR-optimised, and further statistical learning was applied to the qPCR dataset to identify the optimal classifier, which was locked-down for further testing. Independent validation was sought in separate cohorts of CD (n=66) and UC patients (n=57).

**Results:** In both independent validation cohorts, a 17-gene qPCR-based classifier stratified patients into two distinct subgroups. Irrespective of the underlying diagnosis, IBDhi patients (analogous to the poor prognosis IBD1 subgroup) experienced significantly more aggressive disease than IBDlo patients (analogous to IBD2), with earlier need for treatment escalation and more escalations over time.

**Conclusion:** This is the first validated prognostic biomarker that can predict prognosis in newly-diagnosed IBD patients, and represents a step towards personalised therapy.

**What is already known about this subject?:** The course of CD and UC varies considerably between patients, but reliable prognostic markers are not available in clinical practice. This hinders disease management because treatment approaches that would be optimal for patients with indolent disease – characterised by infrequent flare-ups that can be readily controlled by first-line therapy – will inevitably undertreat those with progressive disease. Conversely, strategies that would appropriately control frequently-relapsing, progressive disease will expose patients with more quiescent disease to the risks and side effects of unnecessary treatment. We have previously described a CD8 T cell gene expression signature that corresponds to differences in T cell exhaustion, is detectable during active untreated disease (including at diagnosis), and predicts disease course in both UC and CD. However, the need for cell separation and microarray-based gene expression analysis would make this difficult to translate to clinical practice.

**What are the new findings?:** We have developed, optimised, and independently validated a whole blood qPCR-based classifier – designed to identify the IBD1 and IBD2 patient subgroups – that can reliably predict prognosis in patients with CD or UC from diagnosis without the need for cell separation. We also present a detailed phenotypic update on the disease course experienced by patients in either the IBD1/IBDhi or IBD2/IBDlo subgroups, incorporating both expanded patient cohorts and substantially longer follow-up. This affords new insights into the spectrum of therapies that are differentially required in these patient subgroups, and reinforces their association with disease prognosis.

**How might it impact on clinical practice in the foreseeable future?:** The qPCR-based classifier has performance characteristics that compare favourably to prognostic biomarkers currently in use in oncology, and should be sufficient to guide therapy from diagnosis in patients with CD or UC. This represents an important step towards personalised therapy in inflammatory bowel disease.

## INTRODUCTION

In recent years, there has been a growing realisation that the future of IBD management needs to incorporate a personalised approach to therapy, in which the right treatment can be given to the right patient at the right time.^1^ This is now recognised as a key goal and was recently named as one of the most important research priorities in IBD by the James Lind Alliance priority-setting partnership – a group of clinicians, patients and other stakeholders – who were tasked with identifying important areas of unmet need.^2^ In truth, this ambition is shared across many different disease areas; motivated by developments in oncology where personalised therapy has been shown to be possible using biomarkers that can accurately predict cancer outcome and response to therapy.^3, 4^ The potential advantages of personalised medicine in IBD are clear. First, this would anticipate the marked variability in disease prognosis that occurs between patients,^5, 6^ and which means that “one-size-fits-all” approaches cannot optimally treat everyone (either because they are ineffective in some or unnecessarily risky in others). Second, it would enable clinicians to better use the growing armamentarium of IBD therapies to improve clinical outcomes.^7^ For example, it is well-recognised that early use of combination therapy (an anti-TNFα monoclonal antibody and an immunomodulator) is one of the most effective treatments in CD,^8^ particularly when used early in the disease course,^9, 10^ but also that indiscriminate use of this strategy would be prohibitively expensive and expose many patients to side-effects of drugs that they do not require. Unfortunately, in IBD – as in most autoimmune and inflammatory diseases – biomarkers that can reliably predict the course of disease from diagnosis are not available in clinical practice, precluding the delivery of personalised therapy.

We have previously reported that hypothesis-free inspection of CD8 T cell gene expression data from patients with active, untreated autoimmune disease can identify thousands of genes whose differential expression defines 2 distinct patient subgroups.^11, 12^ In all of the diseases studied, including CD and UC, these subgroups were clinically indistinguishable at enrolment, but patients within them subsequently experienced contrasting disease courses; characterised by clear differences in the time to first relapse and the number of treatment escalations required over time.^11, 12^ More recent work has shown that this gene signature is due to inter-patient differences in T cell exhaustion:^13^ the phenomenon by which effector T cells progressively lose their ability to respond to target antigens. T cell exhaustion was originally reported as a consequence of chronic viral infection,^14^ but is now recognised to occur in the presence of persistent auto-antigens.^13, 15^ Consistent with being less able to respond to disease-related antigens, patients with more T cell exhaustion had a better prognosis, characterised by a longer time to first flare and fewer flares over time.^13^

Here, we describe how we have developed, optimised and independently validated a whole blood biomarker – designed to identify the IBD1/IBD2 subgroups – that is able to predict the course of UC and CD from diagnosis. Additionally, we present a detailed phenotypic update as to the clinical consequences of being in the IBD1 (exhaustion low) or IBD2 (exhaustion high) subgroups.

## MATERIALS AND METHODS

### Patient recruitment (Cambridge cohort)

Patients with active CD and UC, who were not receiving concomitant corticosteroids, immunomodulators or biologic therapy, were recruited from a specialist IBD clinic at Addenbrooke’s hospital before commencing treatment. All subjects were recruited between 2008-2014 and were aged 18 years or older. Most (86/118) were recruited at the time of diagnosis. All patients were diagnosed with CD or UC based on standard endoscopic, histologic, and radiological criteria, and were treated in accordance with national (British Society of Gastroenterology) and international (European Crohn’s and Colitis Organisation) guidelines using a conventional step-up strategy within the UK National Health Service. Disease activity was assessed by considering symptoms, clinical signs, blood tests (C-reactive protein, haemoglobin, serum albumin), stool markers (faecal calprotectin) and endoscopic assessment where indicated. To be enrolled, patients had to have active disease confirmed by one or more objective marker (raised CRP, raised calprotectin, or endoscopic evidence of active disease) in addition to active symptoms and/or signs (**Table 1**). Clinicians were blinded to the biomarker results. Detailed phenotyping data were collected prospectively. Ethical approval for this work was obtained from the Cambridgeshire Regional Ethics committee (REC08/H0306/21). All participants provided written informed consent.

**Table 1.** Baseline patient characteristics

|  | CD |  |  | UC |  |  |
| --- | --- | --- | --- | --- | --- | --- |
|  | IBD1 (n = 33) | IBD2 (n = 33) | P | IBD1 (n = 24) | IBD2 (n = 28) | P |
| Age (years) | 30.3 (25.3-36.1) | 30.3 (23.2-38.7) | 0.98 | 43.8 (30.9-50.4) | 40.5 (29.1-54.0) | 0.92 |
| Gender (% male) | 14 (42.4%) | 13 (39.4%) | 1.00 | 13 (54.2%) | 13 (46.4%) | 0.78 |
| Current smoker | 10 (28.6%) | 12 (33.3%) | 0.79 | 1 (4.2%) | 0 (0%) | 0.46 |
| Newly diagnosed | 27 (81.8%) | 24 (72.7%) | 0.56 | 15 (62.5%) | 20 (71.4%) | 0.56 |
| Disease duration (years) | 0.0 (0.0-0.0) | 0.0 (0.0-0.0) | 0.78 | 0.0 (0.0-2.2) | 0.0 (0.0-3.6) | 0.76 |
| Haemoglobin (g/l) | 12.5 (11.7-13.3) | 13.1 (11.8-13.6) | 0.63 | 14.0 (12.8-14.4) | 13.0 (12.3-14.6) | 0.26 |
| CRP (mg/l) | 26 (16-39) | 25 (10-59) | 0.60 | 6 (3-23) | 4 (2-21) | 0.26 |
| Albumin (g/l) | 35 (32-37) | 37 (34-39) | 0.14 | 39 (37-41) | 39 (37-41) | 0.96 |
| Disease distribution: |  |  |  |  |  |  |
| CD - L1 (ileal) | 9 (27.3%) | 13 (39.4%) | 0.43 | - | - | - |
| CD - L2 (ileocolonic) | 11 (33.3%) | 9 (27.3%) | 0.79 | - | - | - |
| CD - L3 (colonic) | 13 (39.4%) | 11 (33.3%) | 0.80 | - | - | - |
| CD - L4 (upper GI) | 2 (6.1%) | 3 (9.1%) | 1.00 | - | - | - |
| Perianal | 6 (18.2%) | 3 (9.1%) | 0.48 | - | - | - |
| UC - E1 (proctitis) | - | - | - | 5 (20.8%) | 8 (28.6%) | 0.75 |
| UC - E2 (left-sided) | - | - | - | 9 (37.5%) | 11 (39.3%) | 1.00 |
| UC - E3 (extensive) | - | - | - | 10 (41.7%) | 9 (32.1%) | 0.57 |
| Prospective follow up (years) | 4.9 (3.6-7.4) | 5.3 (4.3-8.3) | 0.24 | 5.6 (3.6-7.1) | 5.5 (2.4-8.4) | 0.54 |

Data shown in parentheses indicate median (interquartile range) for continuous variables or percentages for dichotomous variables. Statistical significance was calculated using a Mann-Whitney test (2-tailed) for continuous variables and Fisher's exact test (2-tailed) for dichotomous variables. Disease distribution was classified according to the Montreal Classification<sup>27</sup>. CRP, C-reactive protein; GI, gastrointestinal.

### Sample preparation

For assessment of CD8 T cell gene expression, a 110ml venous blood sample was taken from patients at enrolment. Peripheral blood mononuclear cells were immediately extracted by density centrifugation and CD8 T cells were positively selected, as described previously.^16^ Following purification, cells were lysed and lysates were stored at -80°C. RNA was subsequently extracted using RNeasy Mini Kits (Qiagen) and quantified using a NanoDrop1000 Spectrophotometer (Thermo Scientific). Of the total blood draw, 2.5mls were collected into a PAXgene Blood RNA tube IVD (PreAnalytix) which was stored at -80°C. Whole blood RNA was subsequently extracted using a PAXgene 96 Blood RNA kit (PreAnalytix) according to the manufacturer’s instructions. RNA was quantified as described above.

### Microarray processing and analysis

Following assessment of RNA quality (2100 Bioanalyzer, Agilent Technologies), 200ng RNA was processed for hybridisation onto Affymetrix Human Gene ST 1.0 microarrays (CD8 T cell samples, n=118) or Affymetrix Human Gene ST 2.0 microarrays (whole blood samples, n=69) according to the manufacturer’s instructions. Raw data were pre-processed (background corrected, normalised, quality-checked, and batch-normalised) using Bioconductor packages (http://www.bioconductor.org/) in R (http://www.r-project.org/): *affy*,^17^ *vsn*,^18^ *arrayQualityMetrics*,^19^ and *sva*^*^20^*^. For CD8 T cell data, unsupervised consensus clustering was performed to identify the IBD1 and IBD2 subgroups, as previously described.^12^ Of note, IBD1/IBD2 status was not included as a covariate in the batch normalisation of whole blood samples to reduce the possibility of downward bias in estimating the generalisation error during leave-one-out-cross-validation.

### Biomarker development

Following pre-processing, a statistical (machine) learning method – logistic regression with an adaptive Elastic-Net penalty^21^ – was applied to the whole blood gene expression data to identify genes that could be used to calculate the probability of an individual belonging to either the IBD1 or IBD2 subgroups. Penalised regression methods are a useful tool to regularise models, and thus control overfitting, during biomarker discovery.^22^ The adaptive Elastic-Net method in particular combines the strengths of the ridge penalty and the adaptively-weighted lasso shrinkage penalty, and has been shown to be able to address the challenges inherent in these data.^21^ These were: high dimensionality (i.e. number of samples is substantially smaller than number of genes), multicollinearity (i.e. expression of many genes is correlated, with the need to avoid selecting multiple correlated genes in the final model), and requirement for a sparse and interpretable model (i.e. need for a limited number of genes in a final classifier in which the contribution of each gene can be interpreted). The initial model was determined using a classic Elastic-Net (implemented in the *gcdnet* package^23^ in R) followed by adaptive Elastic-Net training implemented using equations reported in the original description of the method.^21^ In brief, the optimal classification rule to identify the IBD1/IBD2 subgroups was learned from the whole blood microarray data by defining a large set of different combinations of model hyper-parameters, which were then used to fit a corresponding number of candidate models to the whole blood expression data.

Model selection was performed using the Bayesian Information Criterion (BIC), where the highest BIC corresponds to the best model (**Supplementary Table 1**). BIC was defined as:

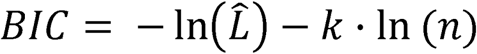

Where k = degrees of freedom (the number of genes incorporated), n = number of samples and (L) = log-likelihood function for the model. The generalisation error of the selected model was estimated using nested leave-one-out cross-validation (LOOCV).^24^

### qPCR classifier development

A list of 39 candidate and 3 reference genes was taken forward to qPCR classifier development using TaqMan gene expression assays (**Supplementary Table 2**). Following reverse transcription of whole blood RNA, qPCR was performed in triplicate using a Roche LightCycler 480, and transcript abundance was calculated using the ΔΔCT method, based on the mean of the technical replicates. The correlation between microarray and qPCR expression values was then used to filter the candidate gene list (6 were removed due to poor correlation). This resulted in a dataset containing expression values for 33 candidate and 3 reference genes from 69 samples. Following normalisation by feature standardisation, an identical penalised regression strategy was applied to this qPCR dataset, to identify an optimal classification model comprising 16 informative and 2 reference genes. To refine this model for use on unscaled data, a pre-requisite for use in a clinical setting, an additional round of penalised logistic regression was applied using the *cvglmnet* function in the *glmnet* package^22^ in R. This uses iterative cross-validation undertaken concurrently to facilitate automatic identification of the optimal, or most regularised, model (using accuracy of IBD1/IBD2 classification as a performance metric). This identified a 17 gene model (15 informative and 2 reference genes) with an error within 1 standard error of the minimum mean cross-validated error, that was considered the most regularised (as recommended by the authors of this approach^22^). This 17 gene classifier (comprising 15 informative and 2 reference genes) was “locked-down” so that no further changes could be made, and was then tested in the validation cohorts. Patients in the qPCR subgroup analogous to IBD1 were termed “IBDhi” and patients in the subgroup analogous to IBD2 were termed “IBDlo”.

### Validation cohorts

One hundred and twenty-three patients with active, untreated IBD (66 CD, 57 UC) were recruited from specialist clinics in 4 UK teaching hospitals (in Cambridge, Nottingham, Exeter and London). Of these patients, 115 (93%) were newly diagnosed (61 CD, 54 UC). Prospective follow-up data were collected for all patients, who were treated at the discretion of their gastroenterologists in accordance with national and international guidelines.

Clinicians were blinded to gene expression analyses. From each patient a 2.5mls venous blood sample was collected into a PAXgene Blood RNA tube IVD (PreAnalytix) which was stored at -80°C. RNA was subsequently extracted, quantified and quality-checked as described above. qPCR was then performed for the 15 informative genes and 2 reference genes within the optimal classifier using RUO (Research-Use-Only) PredictSURE IBD kits (PredictImmune) to determine whether patients were IBDhi or IBDlo. The clinical course experienced by the IBDhi and IBDlo subgroups was then compared using prospectively collected phenotype data. Importantly, the phenotyping collection and analysis was blinded to the classifier designation and vice versa.

### Statistical analysis

Statistical tests performed during microarray analysis or machine learning are described in the relevant sections. Survival analyses for time-to-first-treatment-escalation were performed using a log-rank test. Comparison of the number of treatment escalations was performed using a Mann-Whitney test (2-tailed for CD8 T cell analyses and 1-tailed for validation cohort analyses). Comparison of the clinical and laboratory data in IBD1/IBD2 patients was performed using Fisher’s test for dichotomous variables or Mann-Whitney test for continuous variables (2-tailed). The α value for these analyses was 0.05. All statistical analyses and reporting were performed in accordance with STROBE guidelines.^25^

## RESULTS

### Whole blood classifier development

We have previously reported that a prognostic biomarker based on IBD1/IBD2 subgroup membership would represent a useful clinical tool, given its performance characteristics.^12^ Nonetheless, it is clear that any assay that requires CD8 T cell purification and microarray analysis would be difficult to translate to clinical practice. For this reason, we investigated whether we could identify the same patient subgroups using whole blood, without the need for cell separation (**Figure 1A**). To do this, we first defined a training cohort of 69 patients (39 CD, 30 UC; 35 IBD1, 34 IBD2) for whom we had both CD8 T cell transcriptomic data and a whole blood PAXgene Blood RNA tube – the latter taken at the same time as the CD8 T cell sample. Fifty of these patients were included in our original report of IBD1/IBD2^12^ and 19 were recruited subsequently. RNA was extracted from PAXgene Blood RNA tubes and genome-wide gene expression was measured by microarray (Affymetrix Human Gene ST 2.0 arrays). The resulting raw data was pre-processed to create a normalised dataset that could be used for classifier development (**Materials and Methods**). To identify a whole blood classifier, we used a machine learning method (logistic regression with adaptive Elastic-Net penalisation^21^) to identify models comprising the smallest number of most predictive genes with least redundancy. A series of potential models were produced (**Supplementary Table 1**) of which the optimal model comprised 12 genes and resulted in accurate identification of the IBD1/IBD2 subgroups (P = 1.6 x 10^-7^ for comparison to a ‘dummy’ classifier using a binomial distribution of samples). The generalisation error for this model was estimated using leave-one-out cross-validation (accuracy = 0.81, 95% CI: 0.70-0.90).

**Figure 1.**
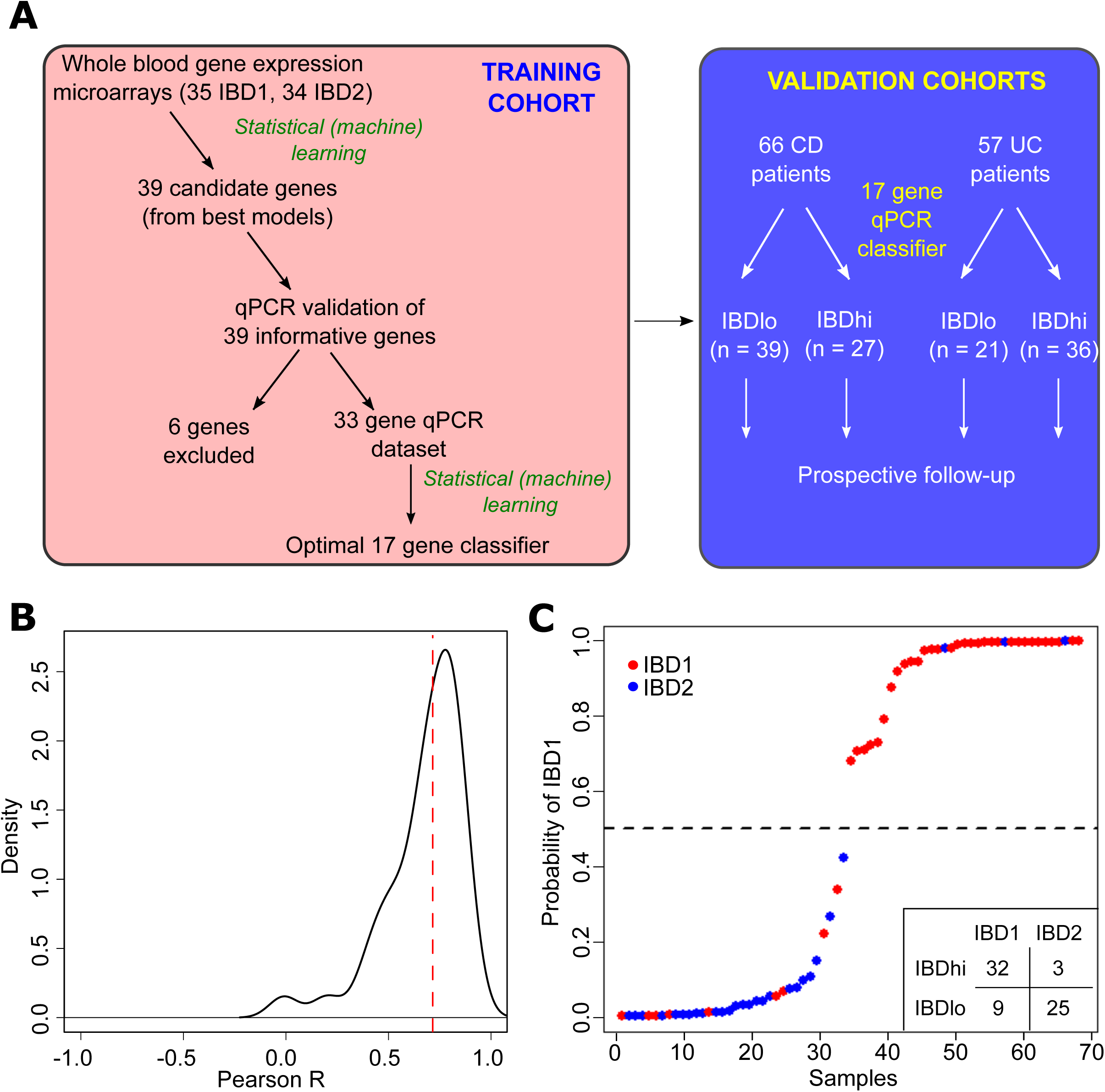
Development of a qPCR-based whole blood prognostic biomarker. **(A)** Schematic depicting the workflow for the development, optimisation and validation of the whole blood qPCR-based classifier with separate training and validation cohorts. **(B)** Distribution of correlation co-efficients between microarray and qPCR-based measurements of gene expression for 39 genes. **(C)** Confidence of assignments to IBD1 and IBD2 subgroups in the training cohort using the qPCR classifier (15 informative and 2 reference genes). Colours indicate actual IBD1/IBD2 assignments based on CD8 T cell transcriptomic analysis (red = IBD1, blue = IBD2). Inset summary table depicts results using 0.5 cut-off for group assignment.

### qPCR classifier development and optimisation

To translate this result into a clinically useful tool, we examined the top models and selected 39 candidate genes and 3 reference genes for qPCR optimisation (**Figure 1A, Supplementary Table 2, Materials and Methods**). Of the candidate genes, 12 were members of the optimal microarray-based classifier, 6 were genes that were highly correlated with genes in the optimal classifier, and 21 were selected from adaptive Elastic-Net models with lower Bayesian Information Criteria (**Supplementary Table 2**). Genes that showed poor correlation with microarray data were excluded (n=6, **Figure 1A-B**). Using qPCR data, we then applied a similar statistical learning strategy (**Materials and Methods**) to identify the optimal classifier (15 informative and 2 reference genes; **Figure 1C, Supplementary Table 3**) which was locked-down for further testing.

### qPCR classifier validation

A critical step in the development of any new biomarker is independent validation, in which the assay can be tested on samples that were not included in the discovery phase. This facilitates an assessment of whether the model will generalise to populations other than the one on which it was developed (**Figure 1A**) and provides a more accurate estimate of the true performance characteristics of the test (or whether the test works at all). We therefore recruited independent cohorts of patients with active, untreated CD (n=66) or UC (n=57) from 4 centres in the UK (Addenbrooke’s Hospital, Cambridge; Royal Devon and Exeter Hospital, Exeter; Nottingham City Hospital, Nottingham; St Mark’s Hospital, London). From each patient 2.5mls blood was taken into a PAXgene Blood RNA tube. Following extraction, quantification and quality checking, RNA was reverse-transcribed and qPCR was performed for the 15 informative and 2 reference genes in the optimal classifier. Using the qPCR data, the classification algorithm assigned every patient into either the “IBDhi” (analogous to IBD1) or “IBDlo” (analogous to IBD2) subgroup. In both the CD and UC validation cohorts, patients in the IBDhi and IBDlo subgroups experienced very different disease courses. Patients in the IBDhi subgroup had consistently more aggressive disease, which was characterised by the need to escalate treatment earlier (with immunomodulators, biologic therapies or surgery) and more frequently than for patients in the IBDlo subgroup (**Figure 2A-F**). In the CD validation cohort, the hazard ratio for the difference in time to first escalation was 2.65 (95% CI: 1.32-5.34; P = 0.006) and in the UC validation cohort this hazard ratio was 3.12 (95% CI: 1.25-7.72; P = 0.015) (**Figure 2A-B**). Moreover, irrespective of the underlying disease, IBDhi patients experienced a disease course that necessitated more potent therapies to achieve disease remission (**Figure 2C-D**). The sensitivity and specificity for predicting the need for multiple escalations within the first 18 months were 72.7% and 73.2% in CD and 100% and 48% in UC. Importantly, because this test would be used at diagnosis, negative prediction (i.e. correctly identifying patients who do not need additional therapy) is critical^26^ – both to improve resource allocation and not miss a “window of opportunity” to optimally treat patients with progressive disease. In these validation cohorts, the negative predictive value (NPV) for predicting multiple escalations within the first 18 months was high: 90.9% in CD and 100% in UC (**Figure 2E-F**). These results are particularly notable given that the classifier was developed to predict IBD1/IBD2 subgroup membership (being directly assessed against this in the training cohort). In the validation cohorts, however, CD8 T cell transcriptomic data – and thus IBD1/IBD2 subgroup membership – was not available, and so the biomarker had to be validated against the difference in prognosis that was observed in the IBD1/IBD2 subgroups. This is one step removed from how the classifier was developed, and so represents a more difficult benchmark, but is ultimately what a prognostic biomarker would need to predict to be clinically useful.

**Figure 2.**
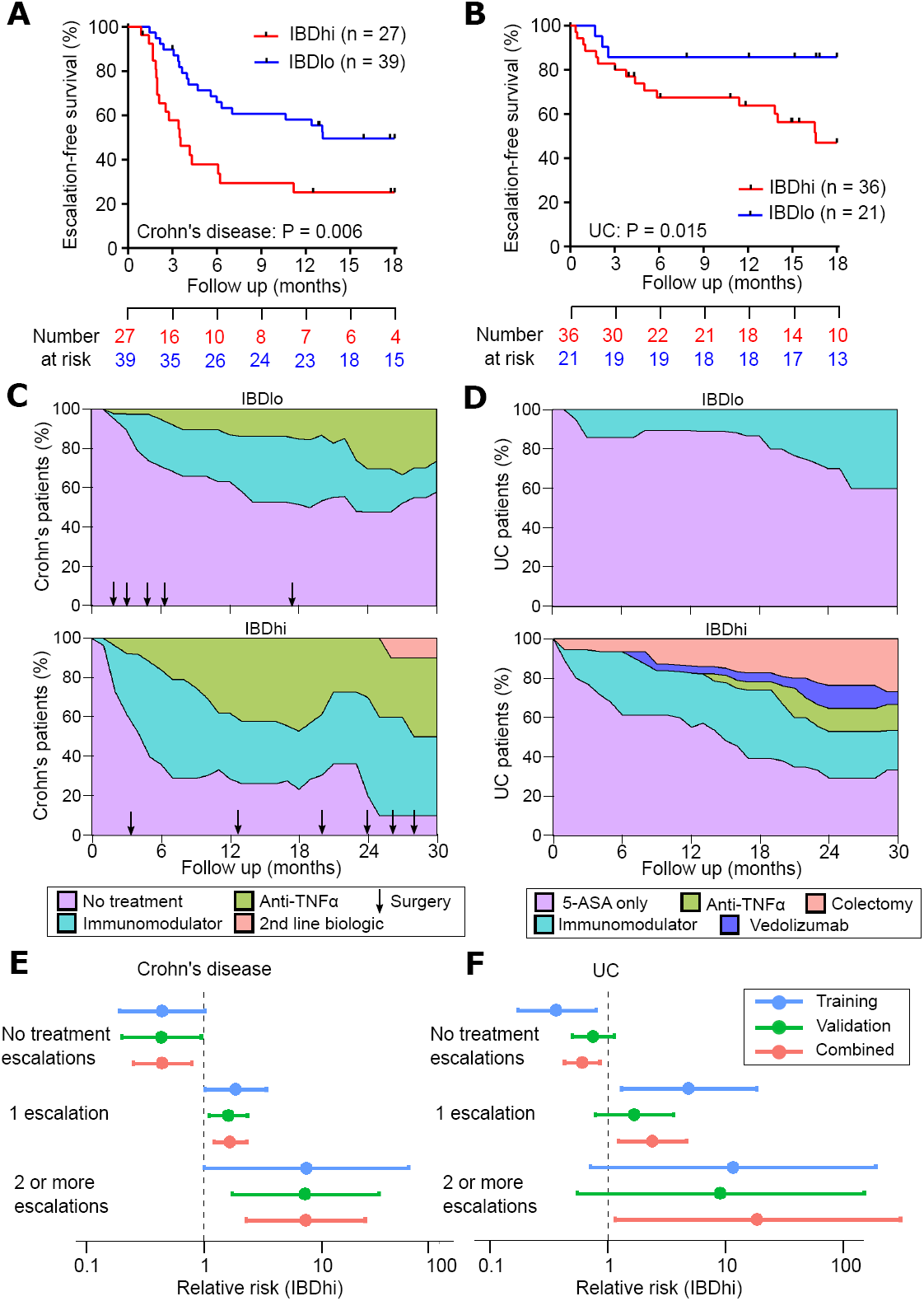
Validation of qPCR-based classifier in independent cohorts. **(A and B)** Kaplan-Meier plots of escalation-free survival for the CD validation cohort (**A**; n = 66) and the UC validation cohort (**B**; n = 57) as stratified by the IBDhi (IBD1 equivalent) and IBDlo (IBD2 equivalent) patient subgroups. Data are censored at 18 months. Statistical significance assessed by log-rank test. **(C and D)** Stacked density plots demonstrating the maximum medical therapy that was required during 2.5 years’ prospective follow up of the IBDhi and IBDlo subgroups in CD **(C)** and UC **(D)**. Treatments were plotted hierarchically (No treatment < Immunomodulator < Anti-TNFα < Second-line biologics [Vedolizumab or Ustekinumab] in CD and 5-ASA only < Immunomodulator < Anti-TNFα < Vedolizumab < Colectomy in UC). Arrows represent episodes of surgery that were required for CD patients at the indicated timepoints. Data are censored accordingly to length of follow-up so that the denominator is the total available cohort at each timepoint. **(E and F)** Forest plots of the relative risk (IBDhi versus IBDlo) of requiring no treatment escalations, 1 treatment escalation or 2 or more treatment escalations within the first 18 months after enrolment for CD patients **(E)** and UC patients **(F)**. Relative risk is with respect to the IBDhi subgroup in each disease, and is presented separately for the training cohort, validation cohort and combined cohorts. Error bars indicate 95% confidence intervals.

To facilitate translation of this test to clinical practice, analytical validation was also performed to formally assess precision, limit of detection, linearity, input RNA range and freeze/thaw cycling for each gene’s qPCR assay and for the combined multianalyte-derived risk score (data not shown). The contribution of specific sources (e.g. operator, batch etc) to the total assay variance was also assessed (data not shown). Together, these analytical and clinical validation data have resulted in a CE-marked assay that is ready for clinical use (PredictSURE IBD, PredictImmune).

### Clinical phenotype over time

It is clear that the phenotypic consequences of IBDhi/IBDlo subgroup membership mirror those previously reported in IBD1/IBD2 patients.^12^ However, due to their prospective collection, both of these cohorts had relatively limited follow up (validation cohort: median 1.7 years; original CD8 T cell cohort manuscript^12^: median 1.6 years). To better understand the longer-term consequences of being in the IBD1 (IBDhi) or IBD2 (IBDlo) subgroups, we examined the extended phenotyping data from all of the patients for whom CD8 T cell gene expression data was available. This cohort was now larger than previously reported^12^ (sample size increased from 67 to 118) and had substantially longer follow-up (median follow-up increased from 1.6 years to 5.3 years). These increases in cohort size and follow-up enabled us to perform a more detailed analysis of the clinical consequences of IBD1/IBD2 subgroup membership. Baseline patient characteristics are presented in **Table 1**. Consistent with our previous findings, all patients could be readily classified into IBD1 or IBD2 based on CD8 T cell gene expression. There were no clinical characteristics at baseline that distinguished between these subgroups (**Table 1**), and in particular there was no correlation between objective measures of inflammation and subgroup membership.

### Disease course in IBD1/IBD2 patients

*CD*: Sixty-six patients with CD were recruited of whom 51 (77.3%) were newly-diagnosed at enrolment. Thirty-three patients were in IBD1 and 33 in IBD2. Compared to patients in the IBD2 subgroup, IBD1 patients had a significantly shorter time to requiring a treatment escalation, as previously reported^12^ (**Figure 3A**). Interestingly, neither clinical parameters (any 2 of: need for steroids, age < 40 years and perianal disease) nor severe endoscopic features at diagnosis (including deep and extensive ulceration in at least one colonic segment) were able to predict the need for early treatment escalation (**Figure 3B-C**). Indeed, even if we attempted to incorporate these, or other, clinical features into a predictive classifier (using a Cox proportional hazards model) we found that none of them were able to improve the performance of the transcriptional classifier (data not shown).

**Figure 3.**
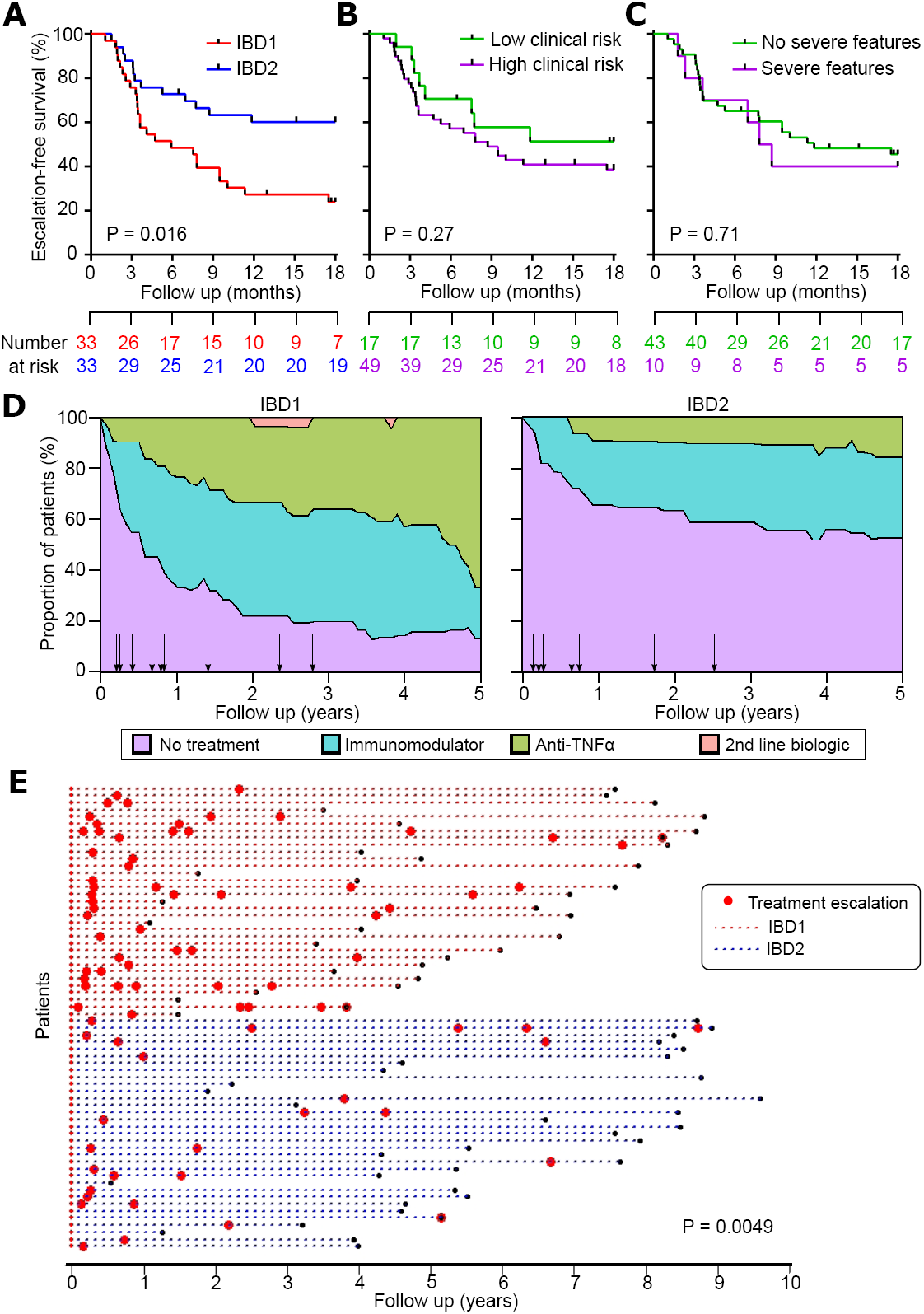
The clinical course of Crohn’s disease is different in IBD1 and IBD2 patients. **(A)** Kaplan-Meier plot of escalation-free survival for CD patients in the IBD1 and IBD2 subgroups. Data are censored at 18 months. Statistical significance assessed by log-rank test. **(B and C)** Kaplan-Meier plots in the same format as (A) with patients subdivided according to clinical risk (high risk = 2 or more of age < 40y at diagnosis, early need for steroids, and perianal disease; **B)** and presence of severe features at index endoscopy (deep and extensive ulceration in at least one colonic segment or endoscopist’s global assessment; **C)**. **(D)** Stacked density plots demonstrating the maximum medical therapy that was required during 5 years’ prospective follow up in the IBD1 and IBD2 subgroups. Treatments were plotted hierarchically (No treatment < Immunomodulator < Anti-TNFα < Second-line biologics [Vedolizumab or Ustekinumab]). Arrows represent episodes of surgery that were required at the indicated timepoints. Data are censored accordingly to length of follow up so that the denominator is the total available cohort at each timepoint. **(E)** Disease course of individual CD patients (dotted lines). The colour of dotted lines reflects subgroup designation. Statistical significance was determined using a Mann-Whitney test.

IBD1 patients with CD also required significantly more treatment escalations over time due to persistently-relapsing or chronically-active disease (**Figure 3D-E**). Indeed, in IBD1 the relative risk (RR) of requiring escalation to biologic therapy (excluding those who received biologic therapy due to immunomodulator intolerance) was 3.0 (12/33 IBD1 patients, 4/33 IBD2 patients) (**Supplementary Table 4**). Likewise, the RR of not requiring any medical therapy in IBD1 was 0.53 (8/33 IBD1 patients, 15/33 IBD2 patients) (**Figure 3E, Supplementary Table 4**). Total surgery rates were similar in both groups (10/33 IBD1, 7/33 IBD2), but all of the patients who required a panproctocolectomy were in the IBD1 subgroup (**Supplementary Table 4**). There were two deaths during follow-up. An IBD2 patient died from end-stage COPD, and an IBD1 patient died from liver failure secondary to PSC.

*UC:* Fifty-two patients with UC were recruited, of whom 35 (67.3%) were newly-diagnosed. Twenty-four patients were in IBD1 and 28 in IBD2. As was observed in the CD cohort, patients in the IBD1 subgroup experienced more aggressive disease with significantly earlier need for treatment escalation (**Figure 4A**). Notably, endoscopic severity at baseline did not predict early need for treatment escalation (**Figure 4B**). Over time, IBD1 patients also required significantly more escalations due to refractory disease (frequently-relapsing or chronically-active) (**Figure 4C-D**). There were several other similarities between the UC and CD cohorts, with the chance of not needing any treatment escalations in IBD1 UC patients being approximately half that of IBD2 UC patients (RR=0.45), and the RR of requiring escalation to biologic therapy or colectomy in IBD1 being 4.08 (7/24 IBD1 patients, 2/28 IBD2 patients) (**Figure 4C-D, Supplementary Table 4**). Indeed, across all of the patient cohorts (CD8 T cell and whole blood) colectomies were only required in IBD1/IBDhi patients (7/56 IBD1 or IBDhi patients; 0/48 IBD2 or IBDlo patients, P = 0.01, two-tailed Fisher’s exact test). There was one death during follow up: an IBD1 patient who was due to start anti-TNFα therapy for chronically-active disease died from a pulmonary embolism during a UC flare.

**Figure 4.**
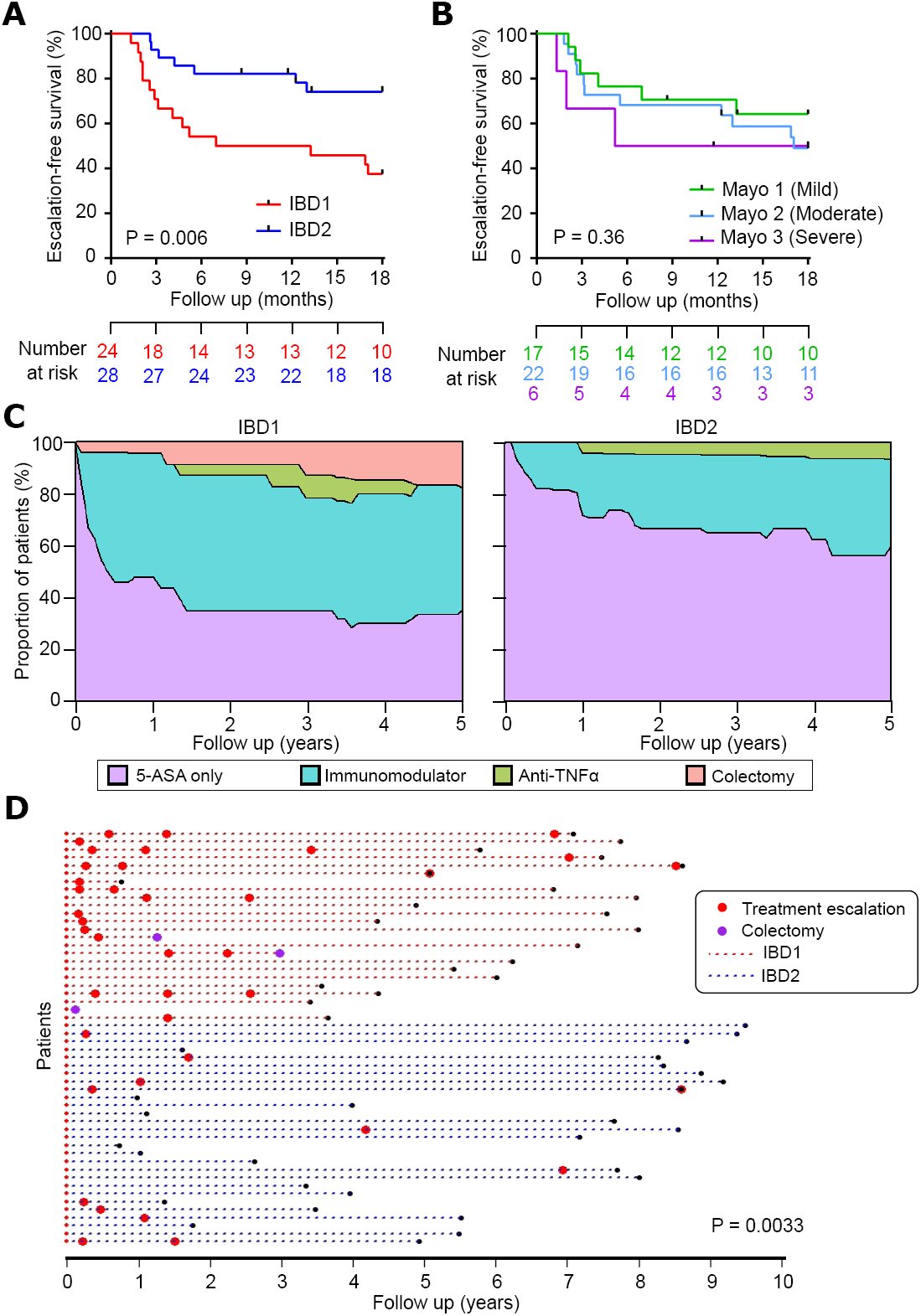
The clinical course of ulcerative colitis is different in IBD1 and IBD2 patients. **(A)** Kaplan-Meier plot of escalation-free survival for UC patients in the IBD1 and IBD2 subgroups. Data are censored at 18 months. Statistical significance assessed by log-rank test. **(B)** Kaplan-Meier plot in the same format as (A) with patients subdivided according to endoscopic disease severity at index colonoscopy.^34^ P value calculated by comparing Mild and Severe cases. **(C)** Stacked density plots demonstrating the maximum medical therapy that was required during 5 years’ prospective follow up in the IBD1 and IBD2 subgroups. Treatments were plotted hierarchically (5-ASA only < Immunomodulator < Anti-TNFα < Vedolizumab < Colectomy). Data are censored accordingly to length of follow up so that the denominator is the total available cohort at each timepoint. **(D)** Disease course of individual UC patients (dotted lines). The colour of dotted lines reflects subgroup designation. Statistical significance was determined using a Mann-Whitney test.

## DISCUSSION

A major barrier to personalised medicine in CD and UC is the lack of suitable biomarkers to guide treatment early in the disease course. Indeed, the requirements for a prognostic test mean that even though several parameters have been associated with prognosis in CD – including clinical features,^28^ serological markers^29^ and genetic variants^30^ – none of these are sufficient to reliably guide therapy in clinical practice. Accordingly, current treatment regimens tend to adopt a "one-size-fits-all" approach, which cannot provide safe, effective and cost-efficient therapy for every patient. Here, we describe the development of a practical, whole blood assay that is in direct response to this unmet clinical need. This assay is the first prognostic test in IBD that has validated performance characteristics that are sufficient to support its use as a prognostic biomarker from diagnosis. Indeed, the performance of the qPCR classifier in both CD and UC is similar to that of existing gene expression-based *in vitro* diagnostic tests in oncology. For example, the hazard ratio for OncotypeDX, a gene expression diagnostic that predicts breast cancer recurrence,^31^ is 2.81 (95% CI: 1.70-4.64).^32^ Importantly, the proven benefit of early aggressive therapy in IBD^9, 10^ should only amplify the clinical benefit of using this assay at diagnosis to stratify patients, since IBDhi patients typically experience the sort of aggressive disease that should benefit most from early use of potent therapy. Collectively, these data support the early adoption of this assay in clinical practice, which should not be logistically difficult since a whole blood qPCR assay can be readily incorporated into standard laboratory protocols.

There are several limitations of this work. First the study was non-interventional and all patients were assessed and treated at the discretion of their gastroenterologists in accordance with national and international guidelines, rather than following a formal protocol. This, however, represents real world practice, and is accordingly the setting in which the test will ultimately be used. Second, while the validated performance characteristics of this assay fulfil the requirements of a useful prognostic biomarker, we have not yet conducted an interventional study to confirm that stratifying therapy using this biomarker would improve clinical outcomes. For this reason, we have concurrently designed a biomarker-stratified trial^33^ to test whether this assay can be used to deliver personalised therapy from diagnosis. This trial (<u>Pr</u>edicting <u>o</u>utcomes for Crohn’s DIsease using a molecular biomark<u>e</u>r; PROFILE; www.crohnsprofiletrial.com) is currently recruiting in the UK, and represents one of the first biomarker-stratified trials in any inflammatory disease. It will assess the relative benefit of “Top Down” therapy (anti-TNFα and an immunomodulator) over “Accelerated Step-Up” therapy in IBDhi and IBDlo patients to determine whether the biomarker can accurately match patients to the most appropriate treatment, thereby improving outcomes by reducing the risk of relapse and minimising drug toxicity.

In summary, we have developed, optimised and validated a whole blood gene expression biomarker that can predict prognosis in patients newly diagnosed with either CD or UC. This provides a rational basis for personalised therapy in IBD, and represents an important step towards precision medicine for patients with CD or UC.

## Acknowledgments

We are grateful to the patients who have provided samples for this study and to the IBD services at Addenbrooke’s Hospital, Royal Devon and Exeter Hospital, Nottingham City Hospital and St Mark’s Hospital for helping identify suitable patients. We also thank Joanne Del Buono, Jennifer Dumbleton, Lawrence Penez, Philip Hendy, Suzie Marriott and Clare Redstone for assistance with phenotyping, and Monica Hou for technical assistance.

## Author contributions

D.B., J.C.L., E.F.M., P.A.L., and K.G.C.S. conceived the study. J.C.L., N.M.N., N.L., T.A., A.H., and M.P. recruited patients and performed prospective phenotyping. D.B. analysed the microarray data to generate a candidate gene list and generated a scaled qPCR-based model. D.R.P. generated qPCR data for use in the final classifier. E.F.M and P.A.L. developed the final model. J.C.L. and K.G.C.S. wrote the final manuscript with input from D.B., M.P., E.F.M., and P.A.L.. All authors reviewed the final manuscript.

